# Silkworm spinning: the programmed self-assembly from natural silk fibroin to superfibre

**DOI:** 10.1101/2021.03.08.434386

**Authors:** Kai Song, Yejing Wang, Wenjie Dong, Zhenzhen Li, Huawei He, Ping Zhu, Qingyou Xia

**Affiliations:** State Key Laboratory of Silkworm Genome Biology, Biological Science Research Center, Southwest University, Beibei, Chongqing, 400715, China; National Laboratory of Biomacromolecules, CAS Center for Excellence in Biomacromolecules, Institute of Biophysics, Chinese Academy of Sciences, Beijing 100101, China; Chongqing Key Laboratory of Sericultural Science, Chongqing Engineering and Technology Research Center for Novel Silk Materials, Chongqing 400715, China; Chongqing Key Laboratory of Soft-Matter Material Chemistry and Function Manufacturing, Chongqing 400715, China; University of Chinese Academy of Sciences, Beijing, 100049, China

## Abstract

Silkworm silk is one of the best natural protein fibers spun by the silkworm at ambient temperature and pressure using aqueous silk protein solution. It is a great challenge to reproduce high-performance artificial fibers comparable to natural silk by bionics for the incomplete understanding of silkworm spinning mechanism, especially the structure and assembly of natural silk fibroin (NSF) in the silk gland. Here, we studied the structure and assembly of NSF with the assistance of amphipol and digitonin. Our results showed NSFs were present as nanofibrils primarily composed of random coils in the silk gland. Metal ions were vital for the formation of NSF nanofibrils. The successive decrease in pH from posterior silk gland (PSG) to anterior silk gland (ASG) resulted in a gradual increase in NSF hydrophobicity. NSF nanofibrils were randomly arranged from PSG to ASG-1, and then self-assembled into herringbone-like patterns near the spinneret (ASG-2) ready for silkworm spinning. Our study reveals the mechanism by which silkworms cleverly utilize metal ions and pH gradient in the silk gland to drive the programmed self-assembly of NSF from disordered nanofibrils to anisotropic liquid crystalline spinning dope (herringbone-like patterns) for silkworm spinning, thus providing novel insights into silkworm/spider spinning mechanism and bionic creation of high-performance fibers.

## Main text

Silkworm silk is one of the best protein fibers in nature by far, which has been utilized by humans for more than 5,000 years as a traditional raw material of textiles. More than 100,000 tons of silk are produced in the world each year (*1*). Sericulture is the main source of family income for millions of farmers in Asia. Today, silk has shown great potentials in flexible electronics, biomedicine and other fields as a new type of material (*2–6*). In addition to silkworm silk, spider dragline silk has also attracted increasing interest, as its comprehensive properties are superior to those of any known synthetic fiber (*2, 7, 8*). However, the commercial era of spider silk has not yet come as spiders are difficult to be domesticated.

It is of great significance to resolve silkworm/spider spinning mechanism for bionics. To date, two models, liquid crystalline spinning (*9*) and micelle models (*10*), have been proposed. The major difference lies in the understanding of silk protein (natural silk fibroin, NSF) structure and assembly *in vivo.* The former suggests that silk protein forms liquid crystalline (*11–13*), while the latter claims that silk protein is present as micelles for its amphiphilic primary sequence (*10, 14, 15*). NSF is stable at 15-30% (w/v) concentrations without precipitation *in vivo* (*2, 16, 17*), whereas it readily transforms from random coil to β-sheet structure *in vitro,* resulting in protein precipitation (*18*). NSF also forms a left-handed 3/2 helix structure specifically at the air-water interface (*19*). Although sericin may protect regenerated silk fibroin (RSF) from aggregation (*20*), it remains unclear how to keep NSF stable *in vitro.* Thus, studying the structure and assembly of NSF *in vivo* is extremely challenging.

Here, we found amphipol and digitonin could keep NSF stable *in vitro* through large-scale screening, and then studied the structure and assembly of NSF *in vivo* using metal shadowing, analytical ultracentrifugation (AUC), fluorescence and circular dichroism (CD) spectroscopy. Our results showed from posterior silk gland (PSG) to anterior silk gland (ASG), NSFs were present as nanofibrils predominantly composed of random coils. Metal ions were crucial for the formation of NSF nanofibrils. NSF nanofibrils were randomly distributed in the lumen from PSG to ASG-1. The hydrophobicity of NSF gradually increased with the decrease of pH from PSG to ASG. Near the spinneret, NSFs self-assembled to form herringbone-like patterns (anisotropic liquid crystalline) ready for silkworm spinning.

## Digitonin/amphipol could stabilize NSF *in vitro*

Large-scale screening showed that membrane scaffold protein MSP1D1 could improve NSF stability from 144 h to 240 h *in vitro* in a concentration-dependent manner. In contrast, BSA decreased NSF stability from 144 h to 96 h (Fig. 1A). MSP1D1 is known as a genetically engineered protein mimicking apolipoprotein A-1 (APOA-1) for the structural study of membrane proteins (*21*). Both APOA and APOE belong to the apolipoprotein family. APOE regulates amyloid-β plaques in the brain of Alzheimer’s patients (*22*). Inspired by the amphiphilicity of MSP1D1, we found amphipol and digitonin were similar to MSP1D1, which could keep NSF stable for 336 h without precipitation (Fig. 1B). Digitonin did not change the secondary structure of NSF (Fig. 1C). Indeed, it stabilized the secondary structure of NSF (Fig. 1D) without altering its state (Fig. 1E). In contrast, NSF transformed rapidly from random coil to β-sheet structure in the absence of digitonin *in vitro* (Fig. 1F), leading to protein precipitation (Fig. 1G). Amphipol was not suitable in CD study for high background noise, although it could also stabilize NSF *in vitro*.

**Fig. 1.**
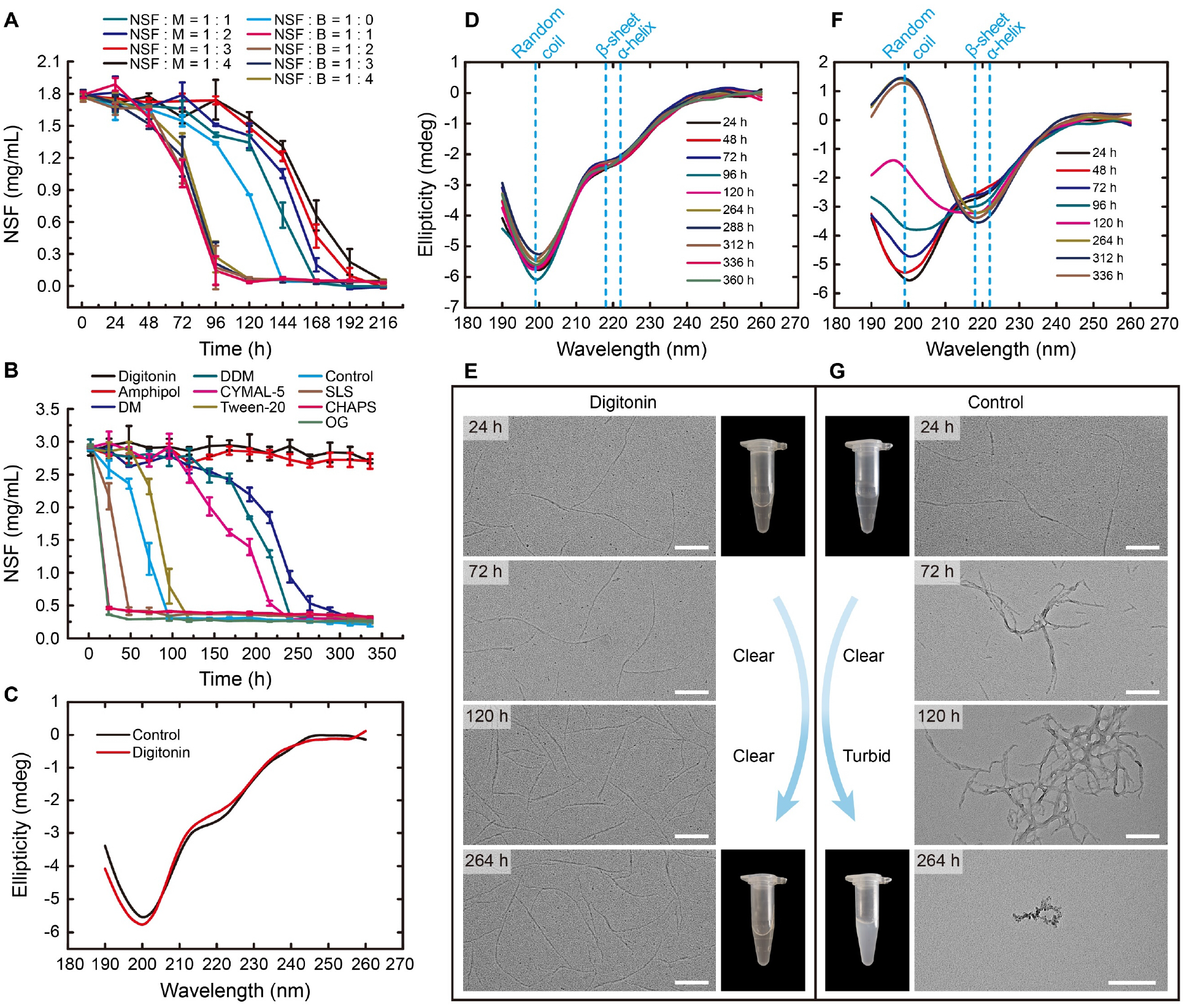
Amphiphilic molecules improved the stability of NSF *in vitro.* **(A)** The changes of UV absorption of NSF (A_280_) over time in the presence of MSP1D1. M, MSP1D1; B, BSA. **(B)** The changes of UV absorption of NSF (A_280_) over time in the presence of different detergents. **(C)** CD spectra of NSF in the presence or absence of digitonin. **(D)** CD spectra of NSF in the presence of digitonin over time. **(E)** The morphology of NSF in the presence of digitonin. The inset showed NSF was stable in the presence of digitonin after 24 h and 264 h, respectively. **(F)** CD spectra of NSF in the absence of digitonin over time. **(G)** The morphology of NSF in the absence of digitonin. The inset showed the aggregation of NSF in the absence of digitonin after 24 h and 264 h, respectively. Scale bar, 200 nm.

## NSFs are present as nanofibrils composed of random coils in the silk gland

With the aid of amphipol/digitonin, we studied the components and properties of NSF. Blue native polyacrylamide gel electrophoresis (BN-PAGE, 3-16%) and SDS-PAGE (4-16%) showed NSF was a macromolecular complex containing four subunits (Fig. 2A), which were further identified as fibroin heavy chain (Fib-H), fibroin light chain (Fib-L), P25 and P25-like by mass spectrometry, respectively (Extended Data Fig. 1).

**Fig. 2.**
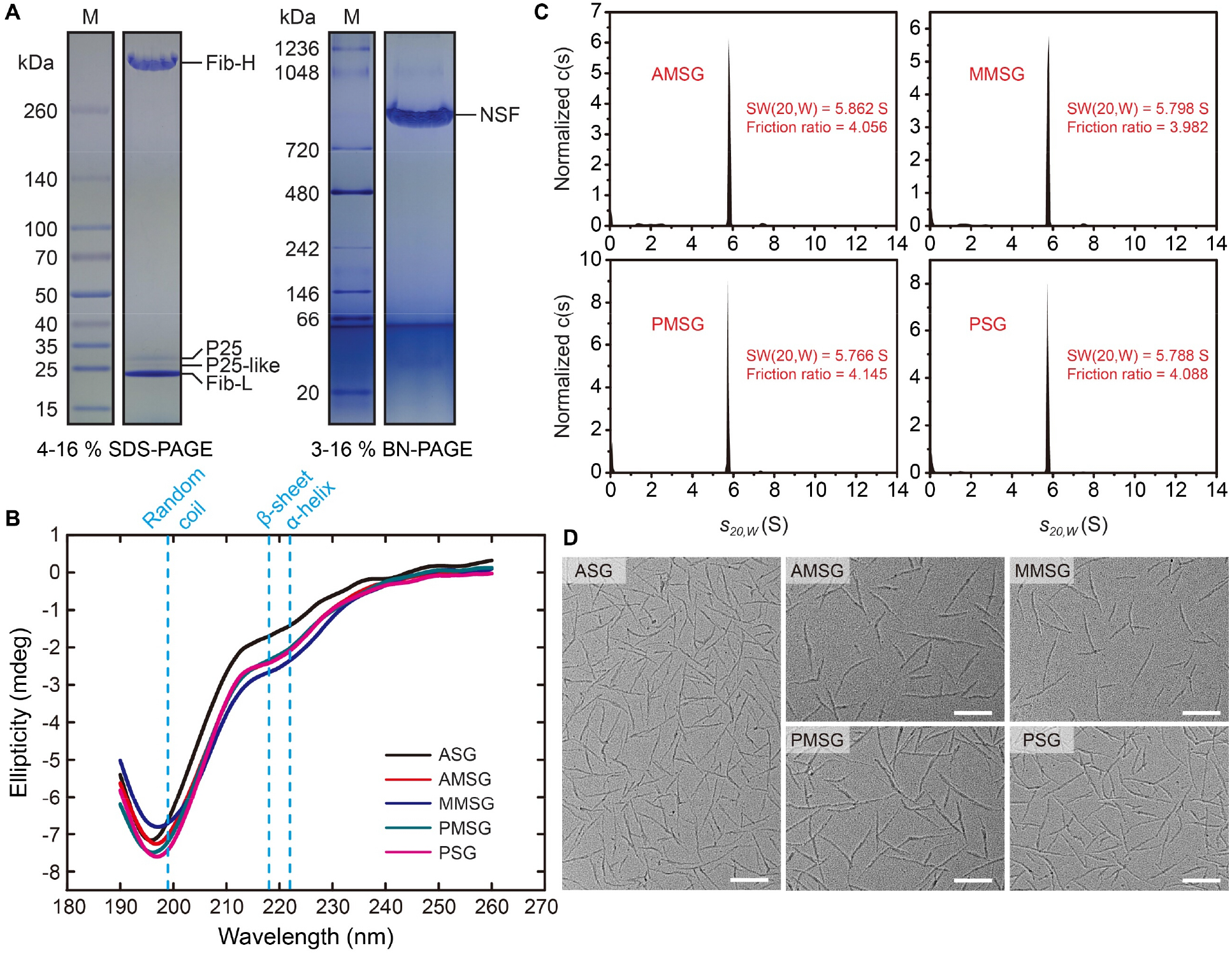
The components and structure of NSF. **(A)** NSF components analysis. NSF was separated by linear gradient SDS-PAGE gel (4-16%) (left) and BN-PAGE gel (3-16%) (right). **(B)** CD spectra of NSF from PSG, PMSG, MMSG, AMSG, ASG. **(C)** AUC analysis of NSF from PSG, PMSG, MMSG and AMSG. The distributions of normalized c(s) were plotted as the function of the sedimentation coefficients *s_20,w_* (S). S, Svedberg (the unit of sedimentation coefficient). **(D)** Metal shadowing of NSF from PSG, PMSG, MMSG, AMSG and ASG. NSF concentration was 0.025 mg·mL^-1^ in 10 mM phosphate buffer. Scale bar, 200 nm.

CD spectra suggested the secondary structure of NSF consisted mainly of random coils and a small number of α-helices from PSG to ASG (Fig. 2B), where NSF from ASG was purified by ions exchange and size exclusion chromatography (Extended Data Fig. 2). RSF was also mainly composed of random coils (Extended Data Fig. 3), which did not change over time under different pH (Extended Data Fig. 4). However, the negative cotton effect of RSF completely disappeared at 222 nm (Extended Data Fig. 3), indicating the difference between NSF and RSF.

AUC showed that the sedimentation coefficients of NSF from PSG, posterior of MSG (PMSG), middle of MSG (MMSG) and anterior of MSG (AMSG) were 5.788 S, 5.766 S, 5.798 S and 5.862 S, and the corresponding friction ratios were 4.088, 4.145, 3.982 and 4.056 (Fig. 2C), respectively. Friction ratios indicated that NSF has a large axial (length-to-width) ratio, implying it may be a fibrous protein. The sedimentation coefficients were very close, indicating NSFs are almost identical from PSG to AMSG.

Metal shadowing showed that NSFs were present as nanofibrils without morphological differences from PSG to ASG (Fig. 2D), which was in line with AUC analysis. Cryogenic transmission electron microscopy (Cryo-TEM) further confirmed the formation of NSF nanofibrils (Extended Data Fig. 5).

## Metal ions induce the formation of NSF nanofibrils

NSF is regarded as a rod-like structure formed by non-covalent aggregation of globular fibroin protein (*23–25*). However, even after incubation with 8 M urea, 2% Triton X-100 and 15 mM dithiothreitol (DTT) for 18 h at 25°C, NSFs were still present as nanofibrils without the appearance of globules (Extended Data Fig. 6, A and B), suggesting NSF are not aggregates of globular protein.

Metal shadowing showed that NSF itself did not form nanofibrils in water, while formed nanofibrils in 50 mM NaCl, which disappeared again after dialysis (Fig. 3A). Hence, we investigated the effects of metal ions on NSF nanofibrils. At 1 mM, Na^+^ and K^+^ induced NSF to form immature fibrous-like structure (Fig. 3B, a and b), while Ca^2+^ and Mg^2+^ induced NSF to assemble into nanofibrils (Fig. 3B, c and d). At 2.5 mM, Na^+^ and K^+^ further induced NSF to assemble into nanofibrils (Fig. 3B, e and f). Once the nanofibrils were formed, they did not change with increasing metal ions concentration, even the concentration increased up to 300 mM (Fig. 3B, g to p).

**Fig. 3.**
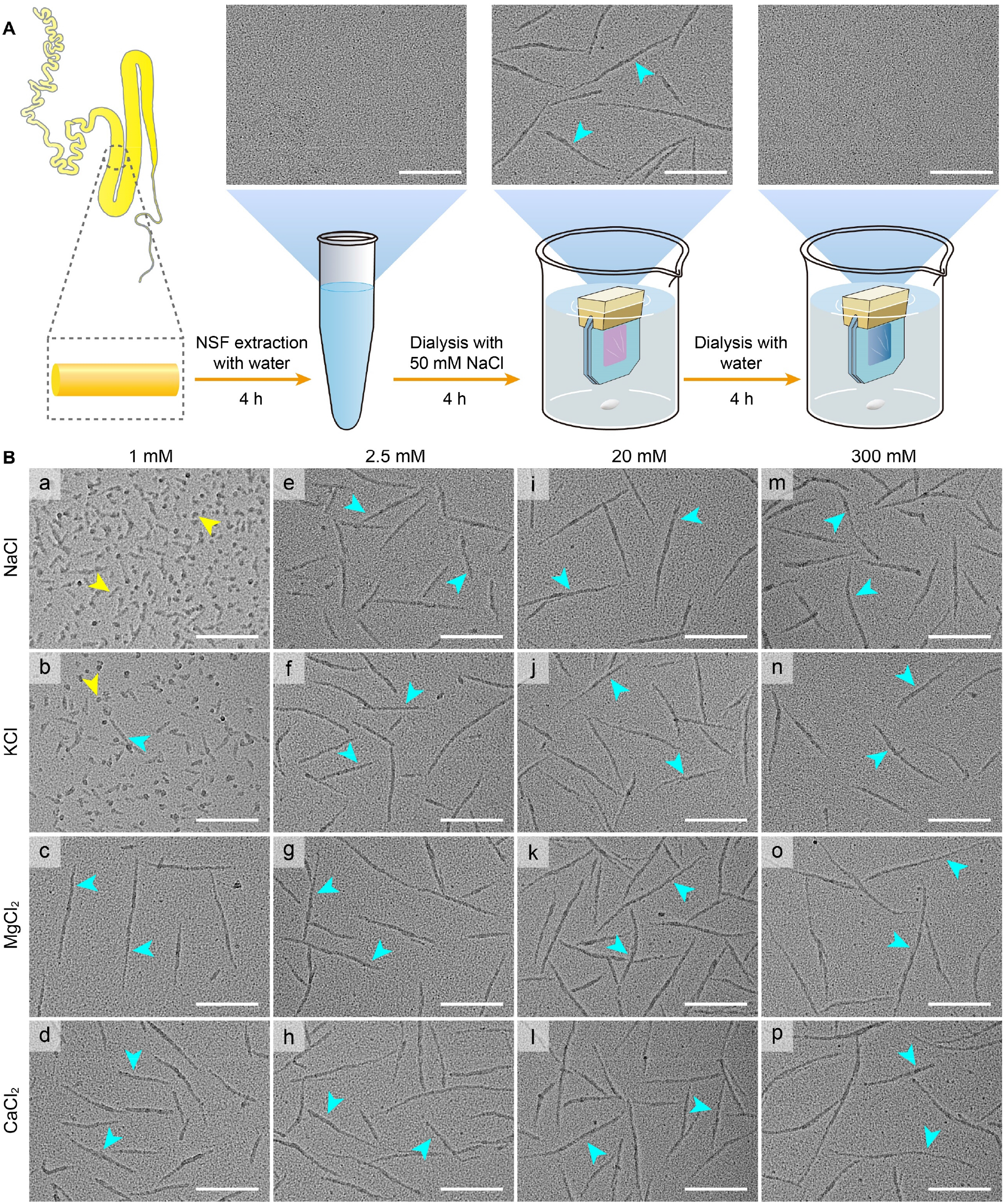
Effect of metal ions on the formation of NSF nanofibrils. **(A)** Metal ions induced the formation of NSF nanofibrils. **(B)** Metal shadowing of NSF in the presence of different concentrations of metal ions. Blue and yellow arrows showed the mature NSF nanofibrils and the immature fibrous-like structure, respectively. Scale bar, 200 nm.

Dynamic light scattering (DLS) showed that the hydrodynamic radius and polydispersion coefficient of NSF were 21 nm and 20.5% in 50 mM NaCl, respectively. After dialysis with water, the hydrodynamic radius slightly decreased to 18.4 nm, and the polydispersion coefficient was multimodal (Extended Data Fig. 6C). No spherical NSF particles with uniform size and smaller hydrodynamic radius appeared after dialysis, suggesting that NSF nanofibrils are not assembled by the aggregation of globular fibroin molecules. The results suggested metal ions are necessary for the formation of NSF nanofibrils. The assembly of NSF nanofibrils induced by metal ions is similar to that of chromatin (*26*).

## The decrease of pH improves NSF hydrophobicity

It is known that the pH in the lumen of silk gland of both silkworms (*27*) and spiders (*28, 29*) continuously decreases from PSG to ASG, which could be mimicked by continuously decreasing pH from 8.0 to 4.8 within 12 h *in vitro* (Fig. 4A). CD spectra showed that pH decreasing did not change the random coil structure of NSF (Fig. 4B), which was consistent with the random coil structure of NSF observed from different segments of the silk gland (Fig. 2B).

**Fig. 4.**
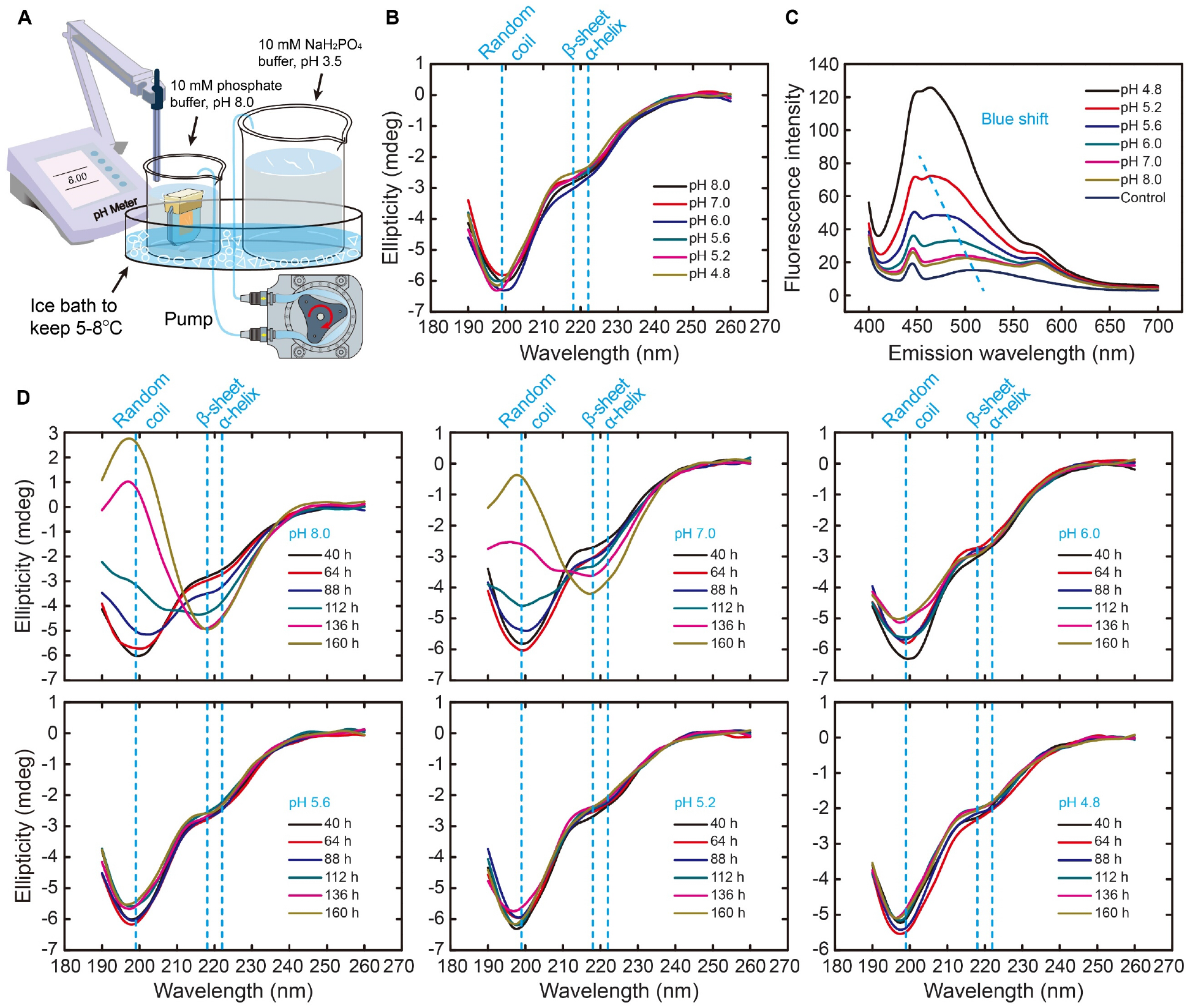
Effect of pH on the structure of NSF. **(A)** Schematic diagram of the formation of a continuous decreasing pH gradient. **(B)** CD spectra of NSF under different pH. **(C)** ANS fluorescence spectra of NSF under different pH. The dotted line showed the blue shift of λmax of NSF. **(D)** Effect of pH on CD spectra of NSF over time.

ANS fluorescence spectra showed that pH decreasing induced a gradual blue-shift of the maximum emission peak (λmax, 507 nm) of NSF (Fig. 4C), indicating a gradual exposure of NSF hydrophobic residues and an increase of NSF hydrophobicity. Similarly, the λmax of RSF gradually blue-shifted with the decrease of pH (Extended Data Fig. 7), indicating pH decreasing improves RSF hydrophobicity, which was in line with the effect of pH on NSF.

The random coil structure of NSF was stable in pH 4.8-5.6 within 160 h, however, it gradually transformed into β-sheet structure in pH 6.0-8.0. Increasing pH promoted the transition of random coil to β-sheet structure of NSF (Fig. 4D). The results suggested that the decrease of pH from PSG to ASG does not change the random coil structure of NSF, but results in the exposure of the hydrophobic residues, thus improving the hydrophobicity of NSF.

## NSFs self-assemble into herringbone-like patterns near the spinneret

It is not yet clear where the liquid crystalline of silk protein is formed in the current liquid crystalline model (*9, 11, 30, 31*). To address this problem, the orientation of NSF was observed in solution and *in situ* in the spinning dope by metal shadowing, respectively (Extended Data Fig. 8). The results showed NSFs were present as nanofibrils randomly arranged in solution. While the concentration was higher than 0.15 mg-mL^-1^, NSF nanofibrils were tightly stacked with random arrangement, and did not change with the increase of NSF concentration (Fig. 5A, a to d).

From PSG to ASG-1 (Fig. 5B), most of NSF nanofibrils were tightly stacked with isotropic orientation (Fig. 5C, a to d), which was consistent with the orientation of NSF nanofibrils in solution (Fig. 5A, c and d). However, a small number of NSF nanofibrils formed an ordered arrangement (Extended Data Fig. 9), which is consistent with the observation of Inoue *et al* (*32*). Interestingly, the long-range ordered molecular alignment of NSF nanofibrils was not observed in the ultra-thin section of ASG-1 (Extended Data Fig. 10), but near the spinneret of ASG-2, where NSF nanofibrils self-assembled into herringbone-like patterns, with the long axes of adjacent molecules aligned parallel to each other (Fig. 5D). The herringbone-like patterns were further packed together to form the spinning dope (Fig. 5E). The results indicated that NSFs are randomly arranged in the lumen from PSG to ASG-1 as isotropic nanofibrils, and self-assemble to form herringbone-like patterns at ASG-2 parallel to the flow direction, which are further packed to form the liquid-crystalline spinning dope with obvious birefringence (*30*).

**Fig. 5.**
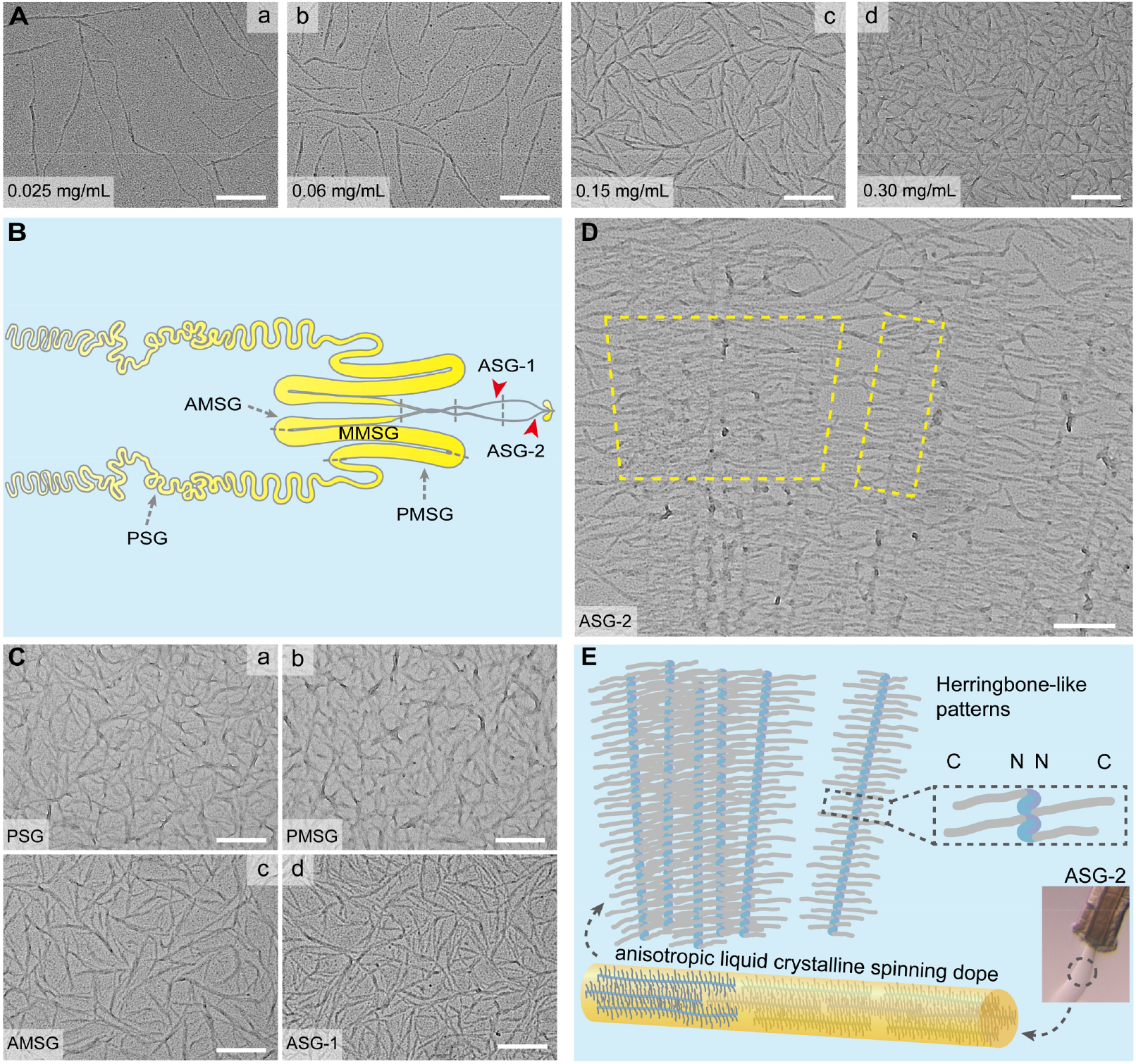
Metal shadowing showed the arrangement and orientation of NSF nanofibrils. **(A)** Metal shadowing of different concentrations of NSF in solution. **(B)** Schematic diagram of different divisions of the silk gland. **(C)** Metal shadowing of NSF *in situ* from different divisions of the silk gland. **(D)** Herringbone-like patterns (anisotropic liquid crystalline phase) self-assembled by NSF nanofibrils at ASG-2. The yellow boxes indicate representative herringbone-like patterns. **(E)** Schematic diagram of anisotropic liquid crystalline spinning dope self-assembled by NSF nanofibrils. C and N, the C and N terminal domain of Fib-H, respectively. Scale bar, 200 nm.

## Discussion

Understanding the structure and assembly of NSF in the silk gland is vital to reveal silkworm natural spinning mechanism. However, it is a great challenge to study the structure and assembly of NSF as it is prone to aggregation *in vitro.* Although RSF is fairly stable *in vitro* (*33*), it has different rheological behaviors (*34, 35*) and structural properties from NSF (Fig. 4, B and D and Extended Data Fig. 3 and 4). Thus, RSF is not an ideal alternative to NSF.

To better study the structure and assembly of NSF, we screened and found digitonin and amphipol could stabilize NSF structure *in vitro* (Fig. 1). Digitonin and amphipol likely interact with NSF through hydrophobic interactions, thus preventing NSF aggregation by shielding the hydrophobic regions of NSF. Although it remains unclear how spiders and silkworms keep high concentrations of silk protein stable *in vivo (16, 18, 36*), our results establish a fundamental basis for studying the structure and assembly of NSF.

The current silk spinning models have shown different understanding on the assembly of silk protein. The micelle model suggests NSFs form micelles (100-200 nm) in solution. With the increase of NSF concentration, micelles coalesce into larger globules (0.8-15 μm), which are further aligned to form fibers under the action of shearing force (*10, 16, 39*). However, the liquid crystalline model indicates the spinning dope in the spider gland and duct forms a nematic phase consisting of rod-like structures, which are essentially aggregates of spherical silk protein (*9, 23*).

Previous evidences show NSF is a large molecular complex (2.3 MDa) composed of Fib-H, Fib-L and P25 as a molar ratio of 6:6:1 (*37*) with a sedimentation coefficient of 10 s (*38*). Here, we identified a new component P25-like in NSF, and demonstrated that NSFs are present as nanofibrils with a sedimentation coefficient of about 5.8 S and a friction ratio of about 4.0 both *in vitro* (Fig. 2, C and D and Extended Data Fig. 3) and *in vivo* (Fig. 5C, a to d). Metal ions are essential for the formation of NSF nanofibrils (Fig. 3A).

Vollrath *et al.* propose the liquid crystalline spinning model (*9*) based on the birefringence of the spinning dope of spiders (*13*) and silkworms (*11*). However, it is not clear whether the liquid crystalline phase is inherent in the silk gland (*9, 11*), or caused by the shearing force during silkworm spinning (*30*). Kerkam *et al.* indicate that liquid crystals are formed after silk protein out of the silk gland but prior to the formation of silk fiber (*31*). Inspired by the discovery of graphene (*40*), we peeled NSF carefully from the spinning dope without altering its orientation, and then determined the molecular orientation of NSF nanofibrils *in situ* in the spinning dope. Metal shadowing showed that NSF nanofibrils are randomly arranged with isotropic orientation from PSG to ASG-1 (Fig. 5C, a to d). Interestingly, NSF nanofibrils self-assemble into herringbone-like patterns with long-range ordered alignment and anisotropic orientation at ASG-2 (Fig. 5, D and E), where silk proteins show obvious birefringence (*30*), indicating NSF nanofibrils form anisotropic liquid crystalline phase in the spinning dope at ASG-2. Our results not only provide direct evidence for the presence of liquid crystalline phase *in vivo*, but also clearly indicate the composition and location of liquid crystalline phase.

The liquid crystalline model suggests NSF forms supramolecular rod-like structures assembled by aggregation of globular silk protein (*23*), however, NSF nanofibrils are very stable without depolymerization into globules even treated by 8 M urea, 2% Triton X-100 and 15 mM DTT at 25°C for 18 h (Extended Data Fig. 6, A and B). DLS indicated NSF nanofibrils may disassemble to form intertwined peptide chains after dialysis with water, rather than globules (Extended Data Fig. 6C).

Our results strongly support that NSFs form nanofibrils instead of the assembly of globules in the silk gland, which are different from the micelles/globules in the micelles model (*10, 16, 39*) and the supramolecular rod-like aggregates of globular silk protein in the liquid crystalline model (*23*). In line with the nanofibrils previously observed in silk fibers (*41–43*), our results provided direct evidence that silk fibers are assembled from nanofibrils.

Although acidification induces the transformation of spider silk protein from random coil to β-sheet structure (*29, 44*), our study suggested that NSFs from different segments of the silk gland are mainly composed of random coil (Fig. 2B), and pH decreasing does not change the random coil structure of NSF (Fig. 4B), indicating acidification (pH decreasing) does not induce the formation of β-sheet structure of NSF *in vivo.* Surprisingly, the β-sheet crystallinity of silk fiber is very small produced by silkworms at low humidity (*45*), implying that NSF itself does not form β-sheet structure *in vivo.* Therefore, the β-sheet structure of silk fiber is most likely formed after NSF is spun out of the silk gland.

Dehydration caused by shearing force and extensional flow is crucial for silk formation (*9, 30*), as sufficiently high concentration of NSF is the basis for silkworms spinning. Here, we found that pH decreasing improves the hydrophobicity of NSF instead of inducing the formation of β-sheet structure (Fig. 4C), which further triggers the separation of NSF from water, thus promoting the dehydration of NSF and an increase of NSF concentration (*9*). Our study suggests that the dehydration caused by pH decreasing is likely crucial for the self-assembly of NSF nanofibrils into herringbone-like patterns (anisotropic liquid crystalline phase) and the final formation of silk fiber.

Obviously, good molecular pre-alignment contributes significantly to the toughness of silk fiber (*9*). Unlike the alignment formed by post-spinning stretching in current artificial spinning, this pre-formed highly ordered molecular alignment may explain the higher toughness of natural silk compared to artificial silk (*46*), and is likely the basis of spider silk as biological superlens (*47*). Unfortunately, it is still unclear how NSF nanofibrils self-assemble into herringbone-like patterns, which is likely initiated by the dimerization of the N-terminal domain of silk protein in response to pH decreasing (*48–50*), and related to NSF concentration (*11, 31*) and shearing force (*30*).

In summary, our results suggest NSFs are present as nanofibrils mainly composed of random coils in the silk gland. Metal ions are indispensable for the formation of NSF nanofibrils. The successive pH decreasing from PSG to ASG improves the hydrophobicity and concentration of NSF rather than inducing the formation of β-sheet structure of NSF. NSF nanofibrils are randomly arranged from PSG to ASG-1, and self-assemble into anisotropic liquid crystalline spinning dope (herringbone-like patterns) at ASG-2 ready for silkworm spinning (Fig. 6). Consequently, silk fibers are formed by the programmed self-assembly of NSF nanofibers undergo an orientation transition from isotropy to anisotropy under the actions of various factors such as decreasing pH gradient, metal ions and shearing force. These findings provide novel insights into silkworm spinning mechanism and biomimetic synthesis of high-performance fibers.

**Fig. 6.**
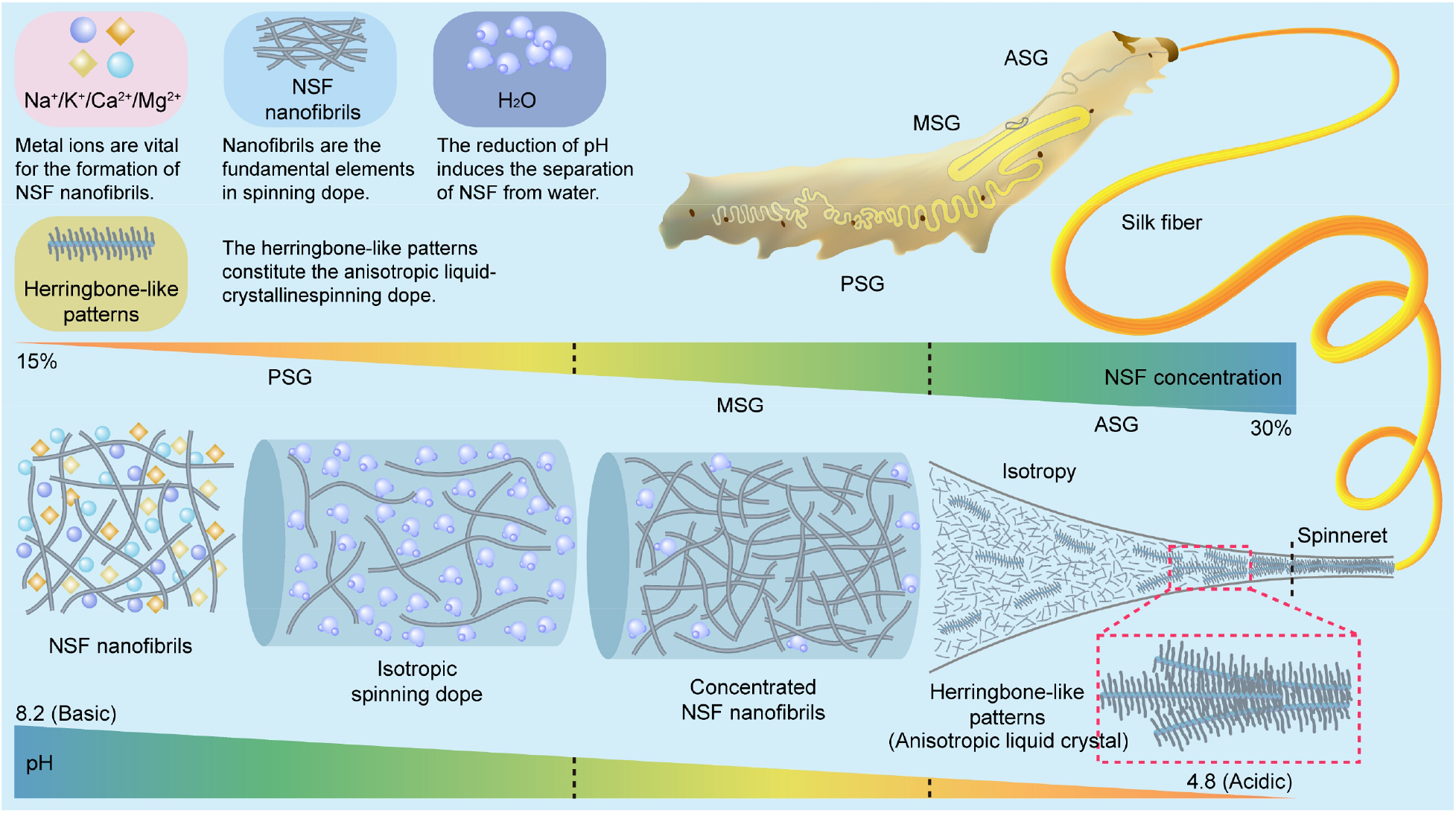
Schematic illustration of the programmed self-assembly of NSF driven by pH gradient and metal ions in the silk gland of the silkworm. **(a)** NSF is a soluble macromolecular complex with a sedimentation coefficient of 5.8 S, and present as nanofibrils induced by metal ions in the silk gland. **(b)** From PSG to ASG, NSF is mainly composed of random coils. With the decrease of pH from about 8.2 in PSG to 4.8 in ASG, the hydrophobicity of NSF is gradually enhanced, and NSF concentration increases up to about 30% (w/v). **(c)** Near the spinneret at ASG-2, NSF nanofibrils self-assemble into herringbone-like patterns (anisotropic liquid-crystalline phase) ready for silkworm spinning.

## Methods

### Isolation and purification of NSF

The silkworm strain Jinsong (*Bombyx mori*), used in this study, was supplied by the State Key Laboratory of Silkworm Genome Biology (Southwest University, Chongqing, China). Silkworm larvae were reared at room temperature until the early wandering stage, then frozen with liquid nitrogen and stored at −80°C. The silk gland is divided into posterior silk gland (PSG), middle silk gland (MSG) and anterior silk gland (ASG) (*51*). Further, MSG has three sections including posterior of MSG (PMSG), middle of MSG (MMSG) and anterior of MSG (AMSG) (*52*). Silk gland was dissected from the frozen silkworm. Silk protein is composed of fibroin and sericin, and stored in the lumen of silk gland (LSG) after synthesis. Sericin could be easily separated from fibroin as it becomes insoluble after freezing. Therefore, natural silk fibroin (NSF) could be purified from the lumen of PSG, PMSG, MMSG and AMSG, respectively. Ultrapure water was prepared by Milli-Q IQ 7000 (Merck, Germany) and used for all tests.

ASG was collected from two hundred fifty living silkworm larvae at the late wandering stage, and then rinsed twice with phosphate-buffered saline (PBS, pH 7.4) for 5 s each. Next, ASG was submerged into 4 mL buffer I (100 mM NaCl, 20 mM Tris-HCl, 0.5 mM EDTA, pH 7.3) precooled at 4°C, and carefully sliced into 3-5 mm segments. After incubation on ice for 4 h, NSF was gradually extracted into buffer I. To keep NSF stable, digitonin was added into the supernatant (0.1%, w/v) after centrifugation at 10,000 g at 4°C for 10 min. Then, the supernatant was incubated with S cation exchange media (200 μL) (Bio-rad, USA) with rotation at 4°C for 2 h. After centrifugation at 1,000 g at 4°C for 10 min, the supernatant was then incubated with Q anion exchange media (200 μL) (Bio-rad) at 4°C for 2 h. Then the media was removed by centrifugation at 10,000 g at 4°C for 10 min. S and Q media were pre-equilibrated with buffer I, respectively. NSF was further purified from the supernatant by size-exclusion chromatography on a Superdex 200 10/300 GL increase column (GE Healthcare, USA) with 10 mM phosphate buffer (Na_2_HPO_4_, NaH_2_PO_4_, pH 8.0) containing 0.015% digitonin. The homogeneity of NSF was assessed by sodium dodecyl sulfate polyacrylamide gel electrophoresis (SDS-PAGE) and Coomassie Brilliant Blue staining (Extended Data Fig. 2). NSF concentration was measured at 280 nm on a NanoDrop 2000C spectrophotometer (Thermo Fisher Scientific, USA). NSF was freshly purified and used immediately to avoid the potential effect of flash-frozen with liquid nitrogen on the structure and properties of NSF.

### Preparation of regenerated silk fibroin

Regenerated silk fibroin (RSF) was prepared as previously described (*33*) with minor modifications. Silkworm cocoons were boiled in 0.02 M Na_2_CO_3_ for 30 min, and then rinsed with water twice to remove soluble sericin from silk fibroin. After drying overnight at room temperature, the degummed silk fiber (silk fibroin) was dissolved in 9 M LiBr (Sangon, China) at 60°C for 20 min to yield 10% (w/v) RSF solution. RSF was dialyzed against water at 4°C for 12 h to remove LiBr, and then dialyzed against 10 mM PBS (pH 8.0) at 4°C for 12 h. After removal of the undissolved aggregates by centrifugation (10,000 g) at 4°C for 15 min, RSF concentration was determined at 280 nm on a NanoDrop 2000C spectrophotometer (Thermo Fisher Scientific).

### Expression and purification of MSP1D1

Membrane scaffold protein MSP1D1 was expressed and purified as described (*53*) with minor modifications. In brief, MSP1D1 was expressed in *E. coli* BL21 (DE3) and induced with 1 mM isopropyl β-D-1-thiogalactopyranoside (IPTG) until the OD_600_ of the cells reached 0.6 at 37°C. After 4 h of induction at 28°C, the cells were collected and then lysed in 20 mM PBS (pH 7.4) containing 1 mM PMSF and 1% Triton X-100. MSP1D1 was purified using his-tag affinity chromatography after centrifugation at 12,000 g for 30 min at 4°C, and then dialyzed against 20 mM PBS (pH 7.4) at 4°C for 24 h to remove imidazole. MSP1D1 concentration was measured at 280 nm using a calculated extinction coefficient of 21,430 L·mol^-1^·cm^-1^. MSP1D1 was concentrated to 5 mg·mL^-1^ and stored at −80°C.

### Polyacrylamide gel electrophoresis

NSF was boiled in reducing buffer for 10 min, and then analyzed by a linear gradient (4-16%) SDS-PAGE gel, which was generated by Hoefer SG 30 gradient maker (Thermo Fisher Scientific). The linear gradient (3-16%) blue native-PAGE (BN-PAGE) gel was cast as witting’s protocol (*54*) using Hoefer SG 30 gradient maker. NSF was incubated with amphipol (Anatrace, USA) at a mass ratio of 1:6 at 4°C for 72 h, then mixed with native-PAGE loading buffer (Invitrogen, USA) and separated by 3-16% BN-PAGE. The molecular weight of NSF was estimated as 445 kDa.

### Circular dichroism

Circular dichroism (CD) spectra were collected from 260 to 190 nm using a 0.1 cm light-path quartz cuvette on a Chirascan Plus spectrometer (Chirascan, UK) with 1 nm bandwidth, 0.5 s response time and 50 nm·min^-1^ scanning speed. NSF was dissolved in 10 mM PBS (pH 8.0) with a final concentration of 0.1 mg·mL^-1^.

### Sedimentation velocity analytical ultracentrifugation

NSF was mixed with amphipol at a mass ratio of 1:6 in buffer II (50 mM NaCl, 50 mM BisTris, 0.5 mM EDTA’ pH 7.5) with a final concentration of 0.75 mg·mL^-1^. Amphipol was dissolved in buffer II with the same concentration as the control. After 72 h of incubation at 4°C, NSF/amphipol (400 μL) and amphipol (400 μL) were loaded into double-sector quartz cells, respectively, mounted into an eight-hole AN-50 Ti rotor, and then centrifuged at 45,000 rpm on ProteomeLab XL-I (Beckman coulter, USA) at 20°C for 5 h. The absorbance at 280 nm was recorded for sedimentation velocity analytical ultracentrifugation (SV-AUC) analysis. The buffer density, viscosity and partial specific volume of NSF were calculated using SEDNTERP (ver. 1.09) to be 1.0018 g·mL^-1^, 1.02298 cP and 0.6945 cm^3^·g^-1^, respectively.

### Metal shadowing

Metal shadowing was performed as Griffith’s method (*55*), as illustrated in figure. S8. For the metal shadowing of NSF in solution, one droplet of NSF (5 μL, 0.025 mg·mL^-1^) was applied to glow-discharged copper grids for 2 min. Excess NSF was carefully removed by blotting paper. For the metal shadowing of NSF *in situ* in the spinning dope, a glow-discharged copper grid was placed on the spinning dope to touch it gently and carefully with tweezers without moving the copper grip to avoid the possible artifact of NSF stacking. NSF was dehydrated in gradient ethanol (0, 25%, 50%, 75%, 100%) for 4 min each, then shadowed with tungsten by DV-502B high vacuum evaporator (Denton Vacuum, USA) followed by air drying. A tungsten wire (8 cm in length) was clamped between two electrodes at a distance of 3.8 cm. The distance of the wire was 9.3 cm from the center of the specimen platform. The angle between the tungsten wire and the sample was about 10°. The total evaporation time was 14.5 min. NSF was kept rotation during the evaporation. Transmission electron microscopy (TEM) was carried out on a Tecnai Spirit (FEI, USA).

### Cryogenic transmission electron microscopy

NSF was purified from PMSG, and then mixed with amphipol in buffer II at a mass ratio of 1:6. The final concentration of NSF was 1.5 mg·mL^-1^. After 72 h of incubation at 4°C, the mixture (3 μL) was loaded on a glow-discharged holey grid (GIG, 1.2-1.3, Au), then vitrified by flash plunging the grid into liquid ethane using vitrobot Mark IV (FEI). The blotting time, force level and humidity were set to be 7 s, 0 and 100%, respectively. Cryogenic transmission electron microscopy (Cryo-TEM) was performed on a 200 kV Talos F200C microscope (FEI) equipped with Ceta camera (FEI).

### Stability analysis of NSF

NSF was purified from PMSG with 20 mM PBS (pH 7.4), and then gently incubated with various detergents at 4°C. The final concentration of NSF and the detergents was 2.93 mg·mL^-1^ and five folds of critical micelle concentration (CMC), respectively, except amphipol was mixed with NSF at a mass ratio of 3:1 (Extended Data Table 1). NSF was incubated with MSP1D1 and BSA as the same procedures. Here, the concentrations of NSF, MSP1D1 and BSA were 2 mg·mL^-1^, 4 mg·mL^-1^ and 10.56 mg·mL^-1^ (Extended Data Table 2 and 3), respectively. Then, the mixture (8 μL) was collected every 24 h, and centrifuged at 10,000 g at 4°C for 10 min. The concentration of the supernatant was determined at 280 nm on a NanoDrop 2000C spectrophotometer (Thermo Fisher Scientific) to value the effects of the detergents, MSP1D1 and BSA on the stability of NSF. Various detergents and BSA were dissolved in water and 20 mM PBS (pH 7.4), respectively, and stored at −20°C.

### Liquid chromatography-tandem mass spectrometry

To identify the components of NSF, NSF was separated by a linear 4-16% SDS-PAGE, and then stained with Coomassie light blue. The target bands were excised for alkylation and reduction with dithiothreitol and iodoacetamide (Sigma, USA), respectively, then digested overnight with trypsin (Promega, USA). Next, the in-gel proteins were extracted by successive washing of gel slices with acetonitrile, and then subjected to liquid chromatography-tandem mass spectrometry (LC-MS/MS) using an LTQ Obitrap XL mass spectrometer (Thermo Fisher Scientific) coupled to a RSLC nano-high performance liquid chromatography (Dionex, USA) system. The peptides were loaded onto a self-packing column (inner diameter, 75 μm; length, 15 cm) filled with 3 μm ReproSil-Pur C18-AQ resin (Dr. Maisch GmbH, Germany), and then eluted with an organic gradient phase (buffer A: 0.5% formic acid/H2O; buffer B: 0.5% formic acid/acetonitrile) at a flow rate of 300 nL/min for 100 min. The program was set as: 0-77 min, 4% buffer B; 78-82 min, 36% buffer B; 83-90 min, 80% buffer B; 91-100 min, 4% buffer B. The datasets were processed with the Proteome Discoverer program (ver 1.4.0.288, Thermo Fischer Scientific). In general, a mass tolerance of 20 ppm for parent ions, 0.6 Da for fragment ions, two missed cleavages, oxidation of Met (dynamic modification) and carbamidomethyl cysteine (fixed modification) were selected as search matching parameters. The results were evaluated using a percolator node (high-confidence q value, FDR < 0.01) to exclude false positives.

### ANS fluorescence spectroscopy

ANS fluorescence spectra were collected on an F7000 spectrophotometer (Hitachi, Japan) in dark at room temperature using a 1 cm light-path cell. NSF (0.3 nM) was incubated with 3 nM ANS in dark under different pH on ice for 3 h. Then, the samples were excited at 388 nm, and the fluorescence emission spectra were recorded from 400 to 700 nm with a scanning speed of 240 nm·min^-1^. ANS was dissolved in water with a stock concentration of 3 mM.

### Ultrathin section of ASG

ASG (including the spinneret) was carefully dissected from mature fifth instar silkworm larvae to ensure that NSF did not leak from the lumen of the silk gland, rinsed twice with 20 mM PBS (pH 7.4) for 3 s each time, and then fixed in 2.5% (v/v) glutaraldehyde/phosphate buffer (0.1 M, pH 7.4) (PB). Next, ASG was fixed with 1% osmium tetraoxide (OsO4) in PB buffer at 4°C for 2 h after rinsed three times with PB buffer, and then dehydrated by gradient ethanol (30%, 50%, 70%, 80%, 90%, 100%, 100%, 7 min each) and acetone twice (10 min each). Subsequently, ASG was sequentially immersed into a graded mixture of acetone and SPI-PON812 resin (19.6 mL SPI-PON812, 6.6 mL DDSA, and 13.8 mL NMA) as the ratio of 3:1, 1:1, 1:3 (v/v), and then infiltrated into the pure resin. Finally, ASG was embedded into the resin with 1.5% BDMA, and then polymerized at 45°C for 12 h and at 60°C for 48 h, respectively. ASG was sliced into ultrathin sections (70 nm thick) by the ultramicrotome EM UC6 (Leica, Germany) using a diamond knife, double stained with uranyl acetate and lead citrate, and then imaged on a Tecnai Spirit transmission electron microscope (FEI).

### Dynamic light scattering

NSF was freshly purified from PMSG with buffer II, and then centrifuged at 10,000 g for 10 min at 4°C to remove the aggregates. The concentration of NSF was 0.5 mg·mL^-1^. Dynamic light scattering (DLS) was performed on NanoStar (WAYTT, USA) to determine the size distribution of NSF before and after dialysis with water. The hydrodynamic radius and the polydispersity coefficient were analyzed using Dynamics (ver 7.1.8.93, USA) and presented as the average from ten independent tests (mean ± SD).

## Data availability

All data are available in the main text or the Extended Data.

## Acknowledgments

We thank Youwang Wang, Wanyu Wu for assistance with the isolation of silk gland, Xiaoxia Yu and Jianhui Li for technical assistance on AUC, DLS and CD spectra, Fuquan Yang, Xing Ding for support with mass spectrometry, Yang Cao for critical reading and helpful discussion. We also thank all members of the center for biological imaging for their support in EM, and in Zhu lab, He lab and Xia lab for their help.

## Author contributions

K. Song and H. He conceived the experiments. K. Song carried out most of the experiments, analyzed the data, composed the figures and wrote the draft. Y. Wang analyzed the data, revised the manuscript and figures. W. Dong performed the ultrathin section. Z. Li analyzed the data. H. He, P. Zhu, and Q. Xia supervised the project, analyzed the data and revised the manuscript and figures.

---

**Extended Data Fig. 1.**
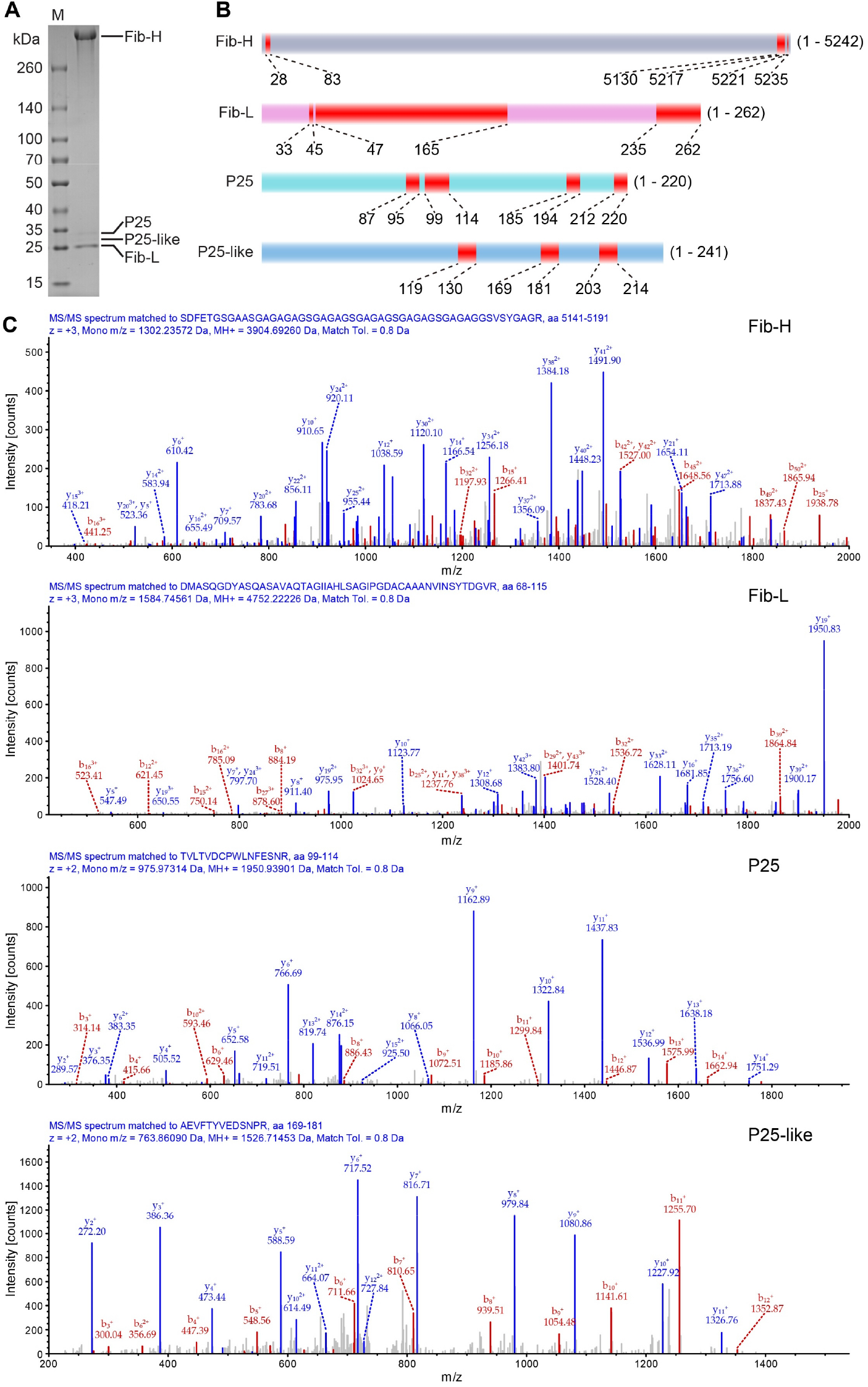
Identification of the subunits of NSF. **(A)** Identification of the subunits of NSF by a linear 4-16% gradient SDS-PAGE. **(B)** Schematic diagram of the subunits Fib-H, Fib-L, P25, P25-like. Amino acid segments identified by LC-MS/MS were indicated in red. **(C)** Typical MS/MS spectra of selected peptides matched to Fib-H, Fib-L, P25, P25-like, respectively. The peaks matched to the peptide sequences were indicated in red (b ion series) and blue (γ ion series), respectively. The unassigned peaks were gray.

**Extended Data Fig. 2.**
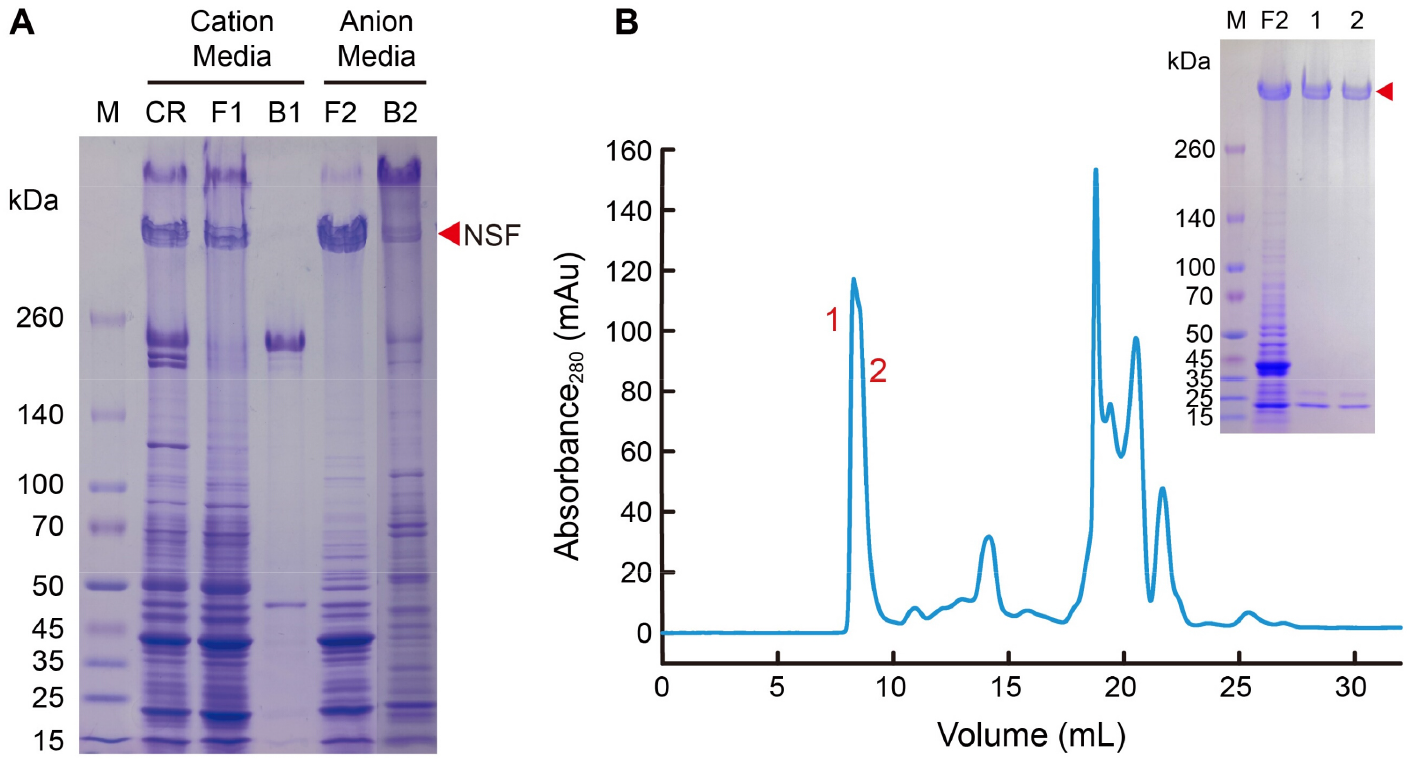
Purification of NSF from ASG. **(A)** A linear 4-16% gradient SDS-PAGE analysis of NSF purified by ions exchange chromatography. M, marker; CR, the crude extraction of ASG; F1, flow-through fraction of the S cation exchange media; B1, beads (S); F2, flow-through fraction of the Q anion exchange media; B2, beads (Q). **(B)** Purification of NSF by size-exclusion chromatography. The inset showed the purity of NSF valued by 4-16% SDS-PAGE. Fractions 1 and 2 were pooled for CD spectra and metal shadowing.

**Extended Data Fig. 3.**
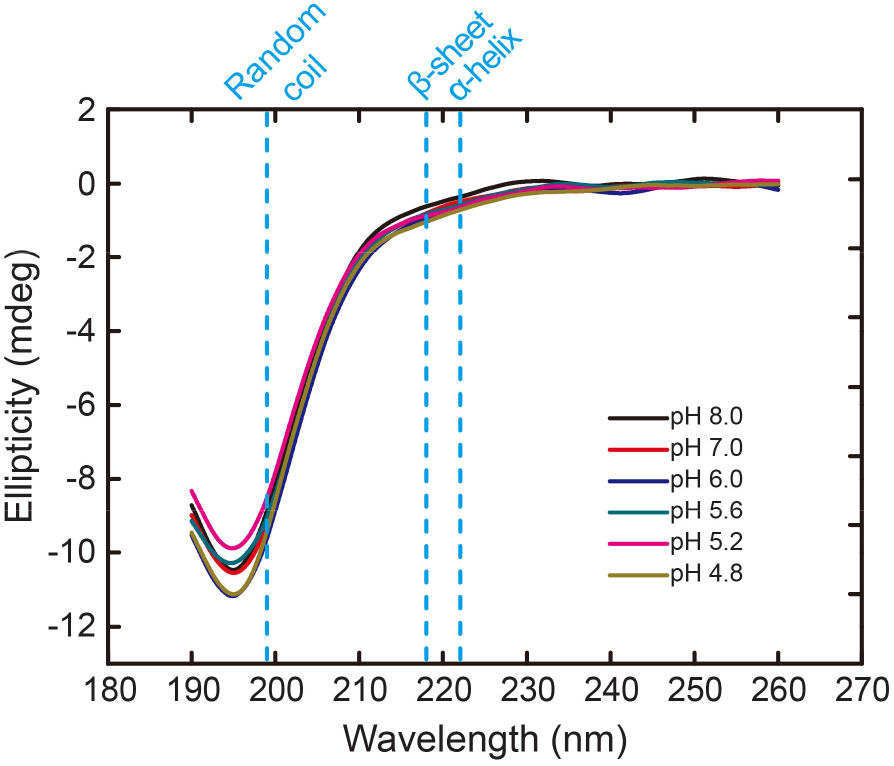
CD spectra of RSF under different pH. The concentration of RSF was 0.1 mg·mL^-1^.

**Extended Data Fig. 4.**
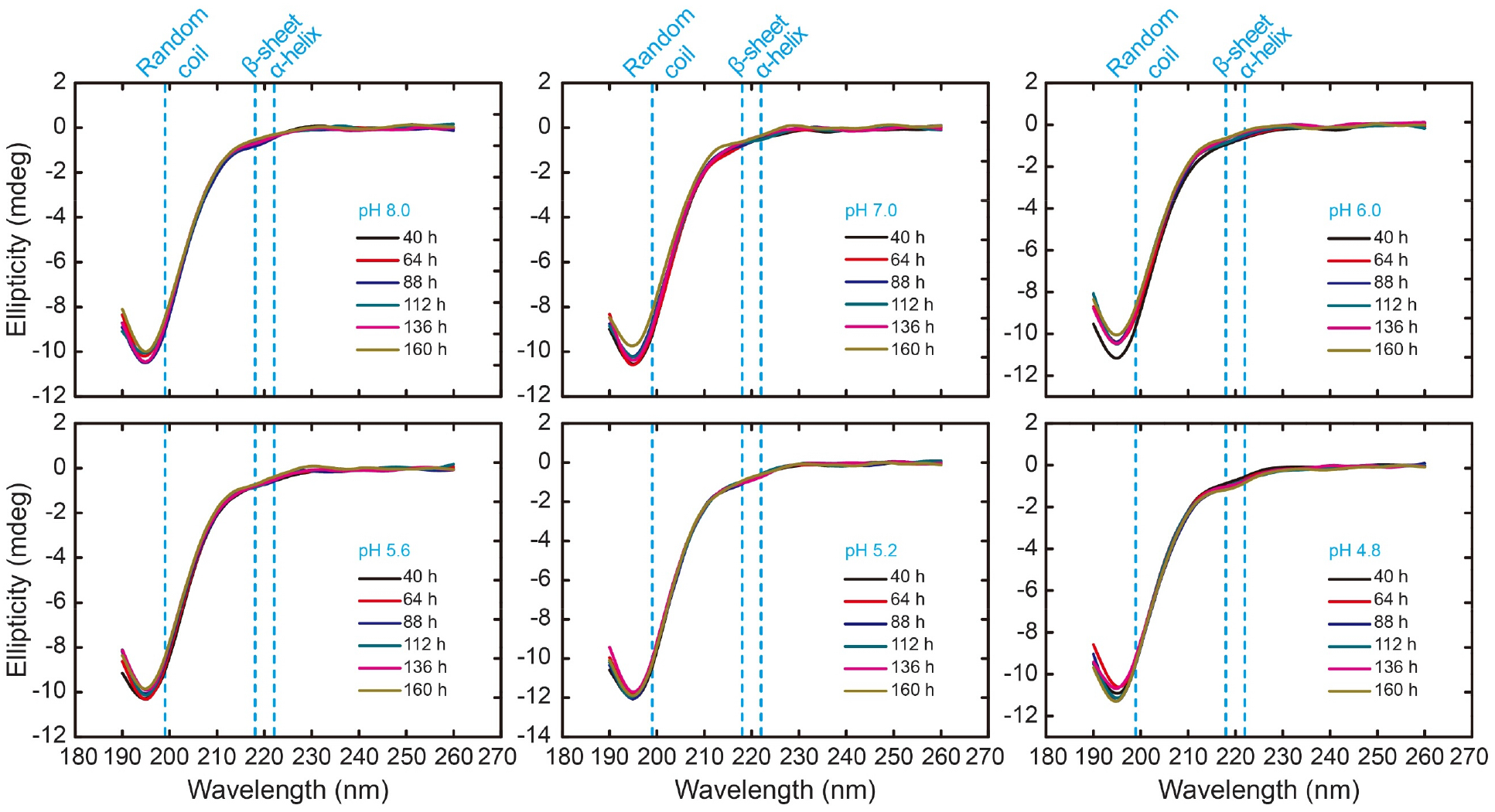
CD spectra of RSF under different pH. The concentration of RSF was 0.1 mg·mL^-1^.

**Extended Data Fig. 5.**
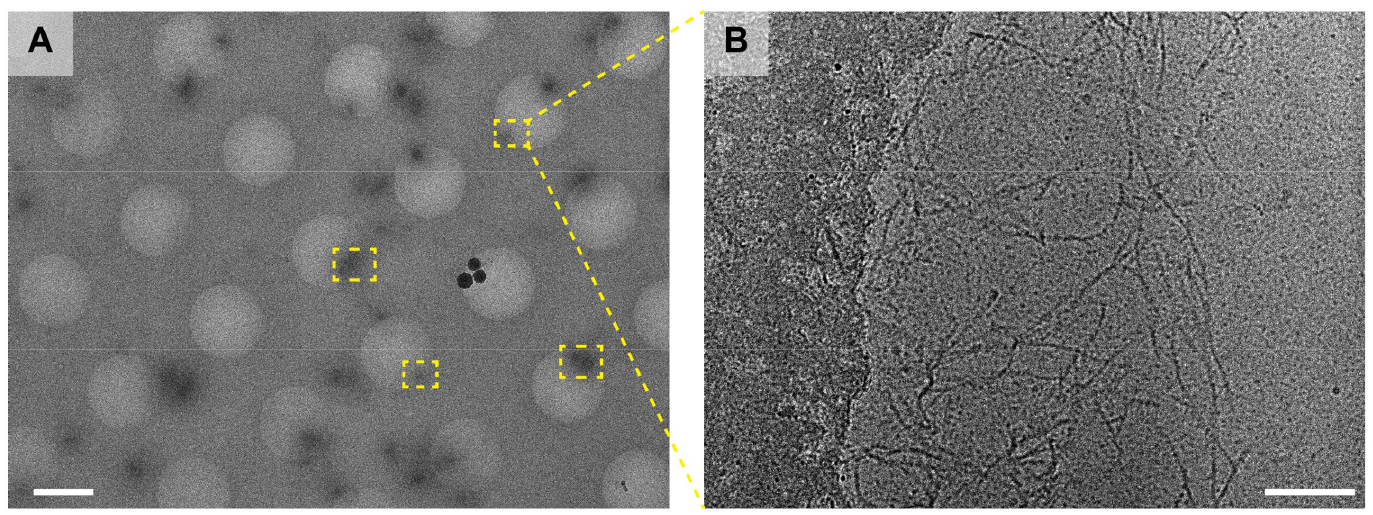
Cryo-TEM of NSF from PMSG. **(A)** Low-magnification image of NSF nanofibrils. Scale bar, 2 μm. **(B)** High-magnification image of NSF nanofibrils. Scale bar, 100 nm.

**Extended Data Fig. 6.**
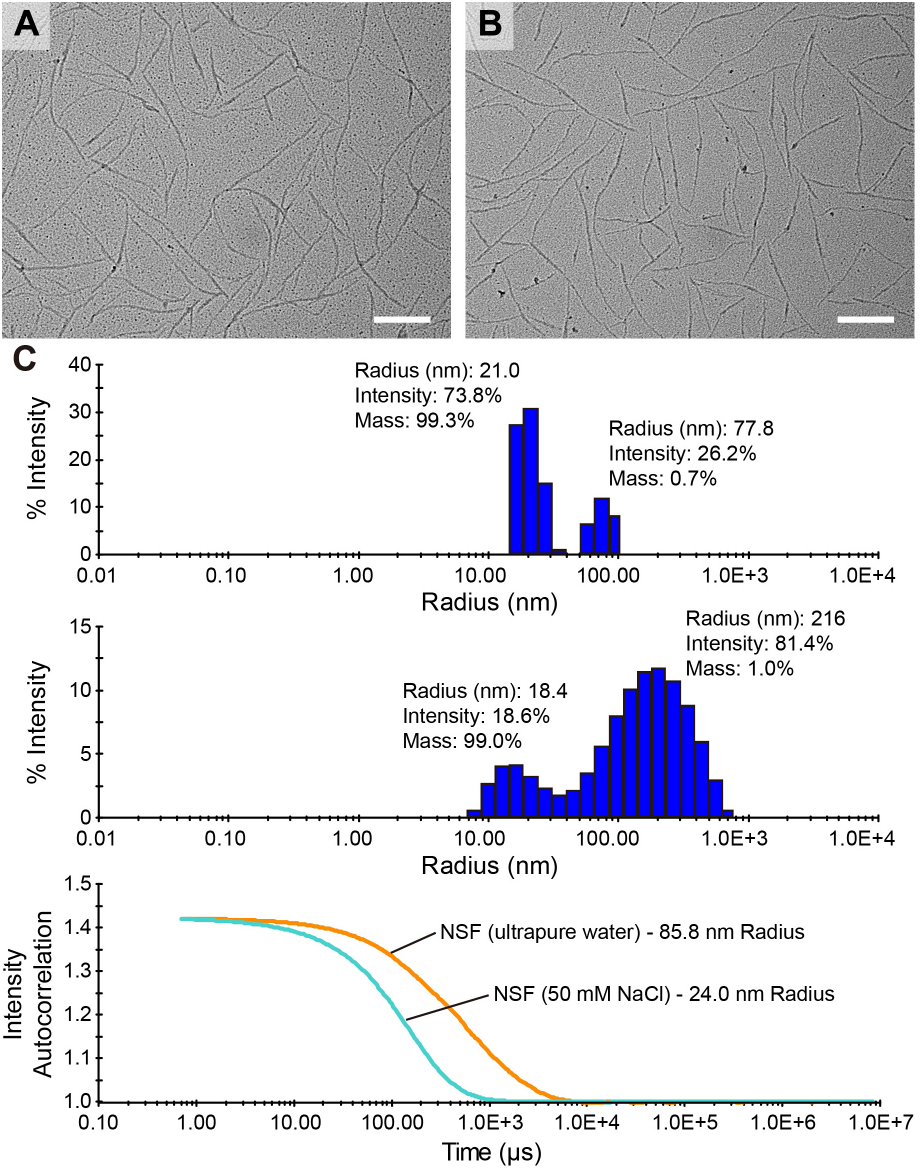
Metal shadowing of NSF. **(A)** Metal shadowing of NSF from PMSG. Scale bar, 200 nm. **(B)** Metal shadowing of NSF from PMSG after 18 h of incubation with 8 M urea, 2% Triton X-100, 15 mM DTT at 25°C. NSF was dissolved in buffer II with a final concentration of 0.025 mg·mL^-1^. Scale bar, 200 nm. **(C)** DLS analysis of the hydrodynamic size distribution of NSF in buffer II (up), after dialysis against water (middle) and the correlation curves (down).

**Extended Data Fig. 7.**
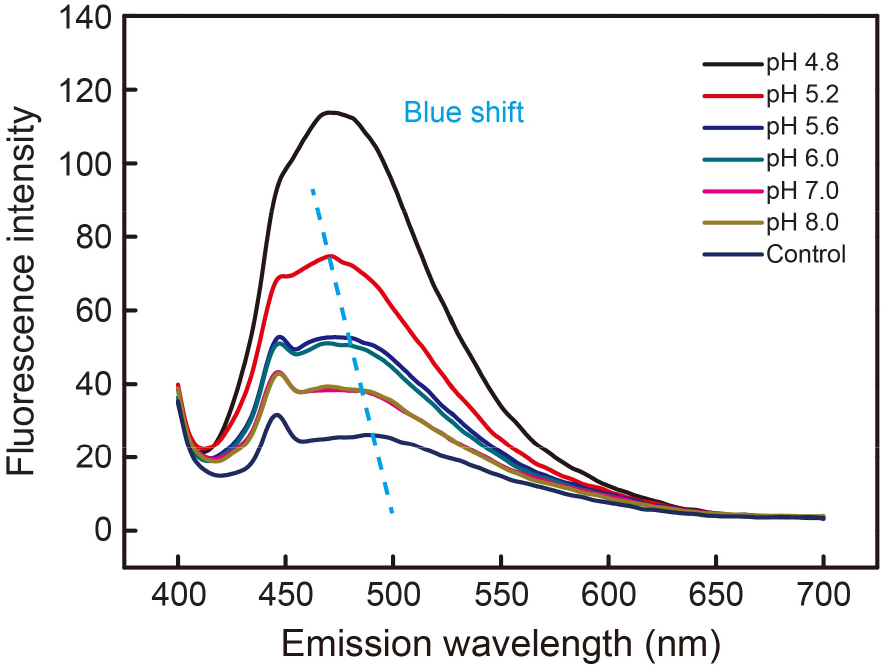
ANS fluorescence spectra of RSF under different pH. The dotted line showed the blue shift of λmax. The concentration of RSF was 0.2 mg·mL^-1^.

**Extended Data Fig. 8.**
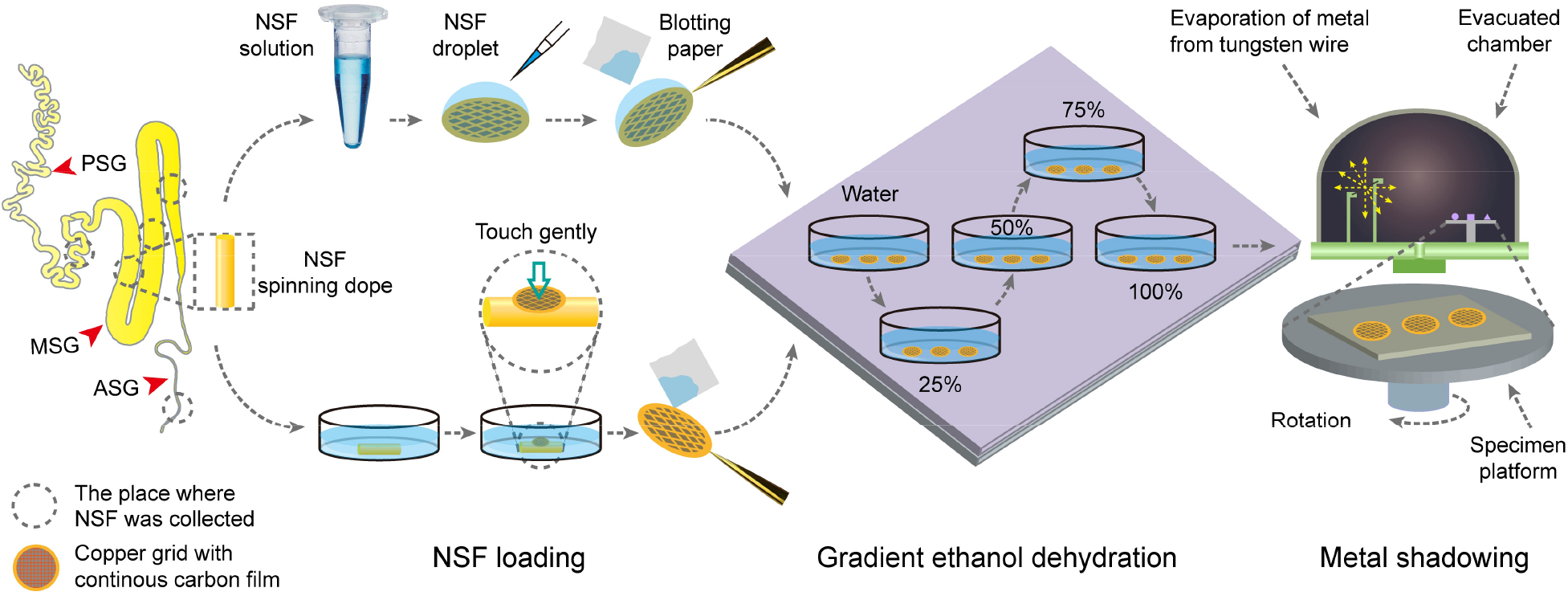
Schematic diagram of the metal shadowing of NSF in water and *in situ* in the spinning dope from the silk gland.

**Extended Data Fig. 9.**
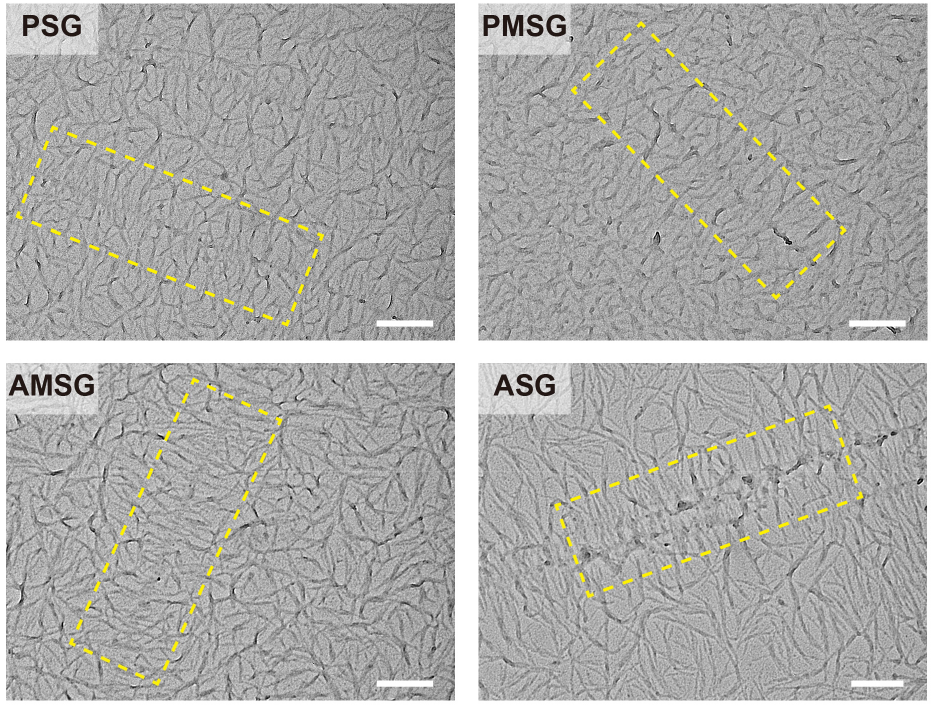
Metal shadowing of NSF *in situ* in the silk gland. Dotted box represented a herringbone-like pattern of NSF nanofibrils. **PSG,** posterior silk gland. **PMSG, AMSG,** the posterior and anterior of middle silk gland, respectively. **ASG,** anterior silk gland. Scale bar, 200 nm.

**Extended Data Fig. 10.**
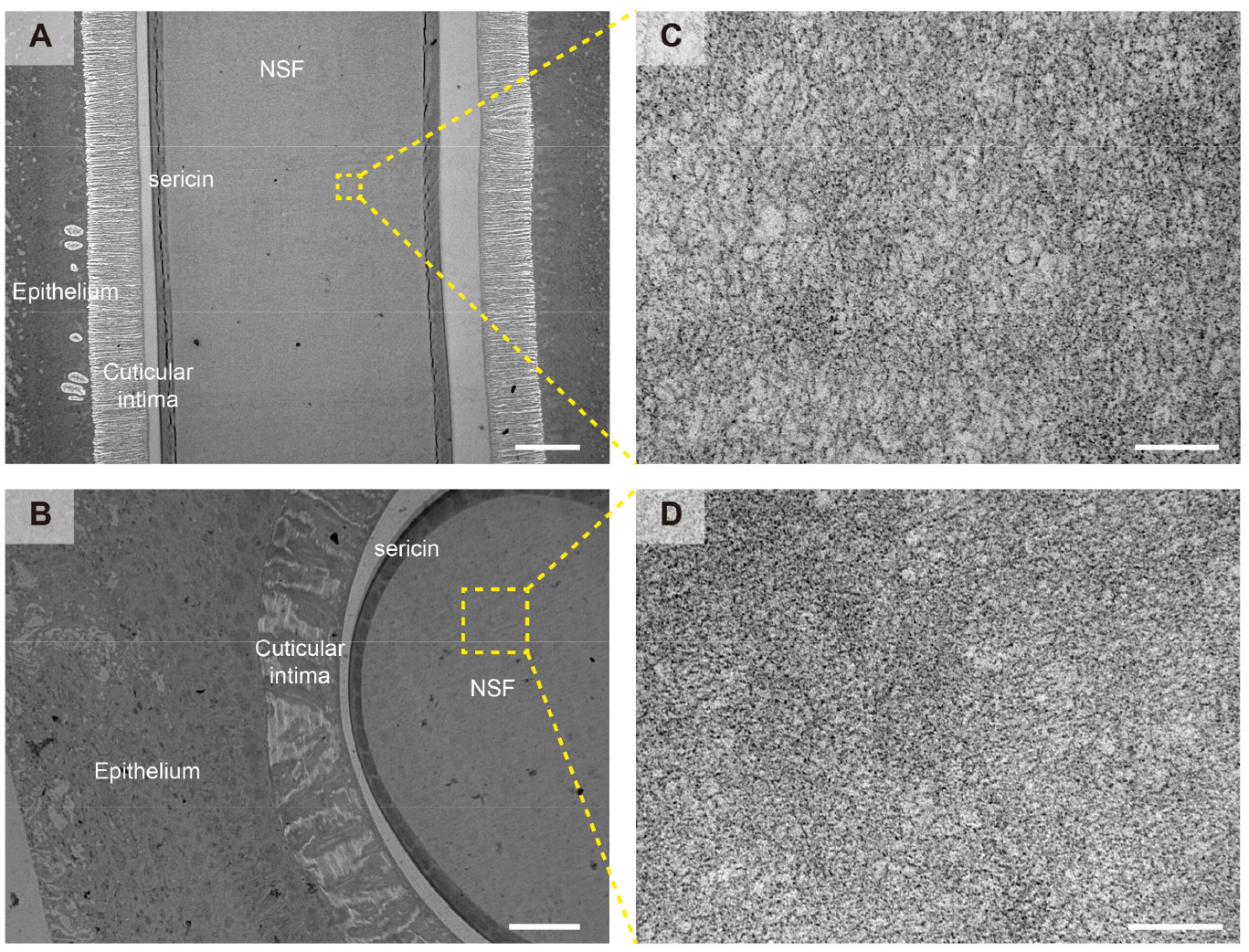
Ultrathin-section TEM of ASG-1. **(A)** Longitudinal section (parallel to the spinning direction) of ASG-1. Scale bar, 50 μm. **(B)** Cross-section (perpendicular to the spinning direction) of ASG-1. Scale bar, 10 μm. **(C-D)** The boxes indicated the location of the magnification, revealing the presence of isotropic NSF spinning dope in the lumen of the silk gland. Scale bar, 200 nm.

**Extended Data Table 1.**
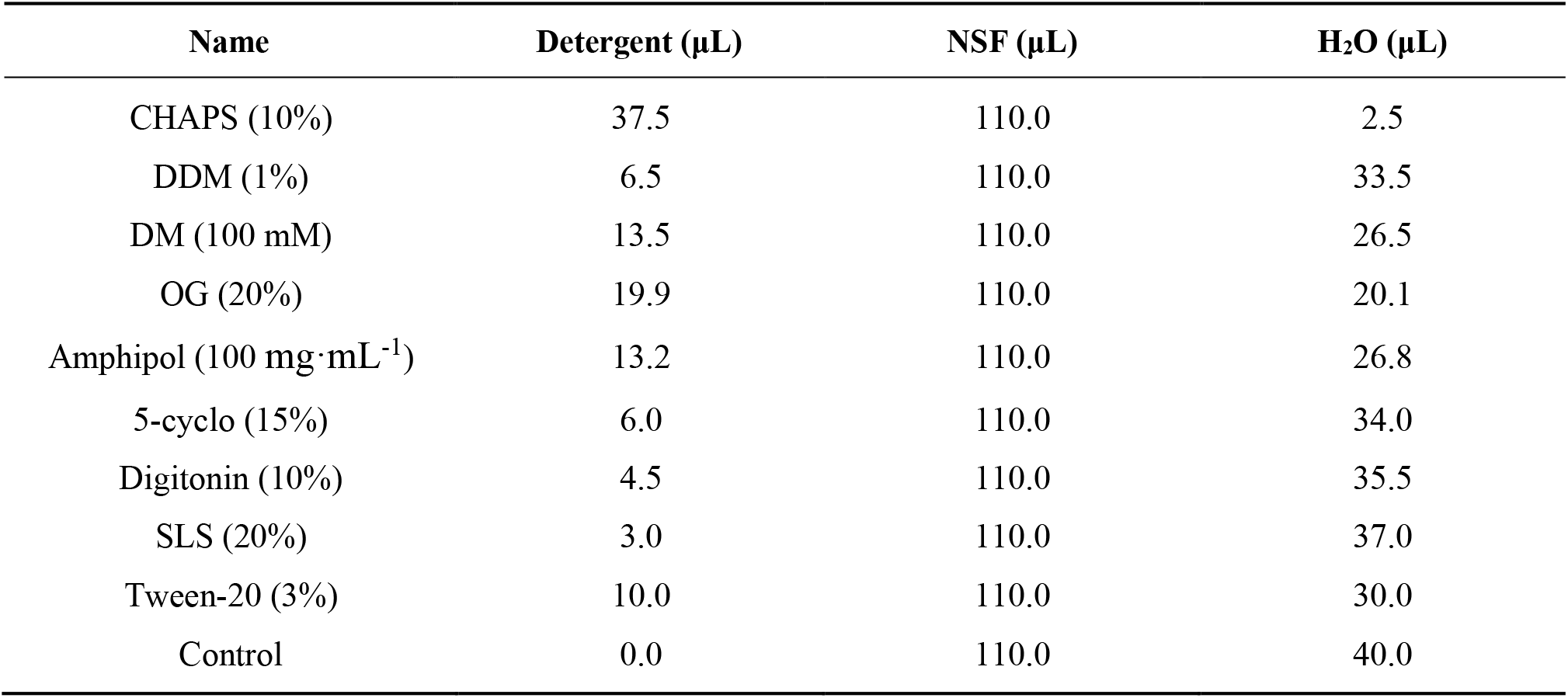
Effect of various detergents on NSF stability

| Name | Detergent ($\mu\text{L}$) | NSF ($\mu\text{L}$) | H<sub>2</sub>O ($\mu\text{L}$) |
| --- | --- | --- | --- |
| CHAPS (10%) | 37.5 | 110.0 | 2.5 |
| DDM (1%) | 6.5 | 110.0 | 33.5 |
| DM (100 mM) | 13.5 | 110.0 | 26.5 |
| OG (20%) | 19.9 | 110.0 | 20.1 |
| Amphipol (100 mg·mL<sup>-1</sup>) | 13.2 | 110.0 | 26.8 |
| 5-cyclo (15%) | 6.0 | 110.0 | 34.0 |
| Digitonin (10%) | 4.5 | 110.0 | 35.5 |
| SLS (20%) | 3.0 | 110.0 | 37.0 |
| Tween-20 (3%) | 10.0 | 110.0 | 30.0 |
| Control | 0.0 | 110.0 | 40.0 |

**Extended Data Table 2.** Effect of MSP1D1 on NSF stability

| **NSF:MSP1D1**<br>**(Molar ratio)** | **NSF**<br>**(μL)** | **MSP1D1**<br>**(μL)** | **PBS**<br>**(μL)** |
| --- | --- | --- | --- |
| Control (1 : 0) | 200.0 | 0.0 | 22.5 |
| 1 : 1 | 200.0 | 5.6 | 16.9 |
| 1 : 2 | 200.0 | 11.2 | 11.2 |
| 1 : 3 | 200.0 | 16.9 | 5.6 |
| 1 : 4 | 200.0 | 22.5 | 0.0 |
| 0 : 1 | 0.0 | 5.6 | 216.9 |
| 0 : 2 | 0.0 | 11.2 | 211.3 |
| 0 : 3 | 0.0 | 16.9 | 205.6 |
| 0 : 4 | 0.0 | 22.5 | 200.0 |

**Extended Data Table 3.** Effect of BSA on NSF stability

| **NSF:BSA**<br>**(Molar ratio)** | **NSF**<br>**(μL)** | **BSA**<br>**(μL)** | **PBS**<br>**(μL)** |
| --- | --- | --- | --- |
| Control (1 : 0) | 200.0 | 0.0 | 22.5 |
| 1 : 1 | 200.0 | 5.6 | 16.9 |
| 1 : 2 | 200.0 | 11.2 | 11.2 |
| 1 : 3 | 200.0 | 16.9 | 5.6 |
| 1 : 4 | 200.0 | 22.5 | 0.0 |
| 0 : 1 | 0.0 | 5.6 | 216.9 |
| 0 : 2 | 0.0 | 11.2 | 211.3 |
| 0 : 3 | 0.0 | 16.9 | 205.6 |
| 0 : 4 | 0.0 | 22.5 | 200.0 |

